# Dynamic RNA binding and unfolding by nonsense-mediated mRNA decay factor UPF2

**DOI:** 10.1101/2024.10.26.620407

**Authors:** Jenn-Yeu A. Szeto, Mirella Vivoli Vega, Justine Mailliot, George Orriss, Lingling Sun, Joshua C. Bufton, Kyle T. Powers, Sathish K.N. Yadav, Imre Berger, Christiane Schaffitzel

## Abstract

Nonsense-mediated mRNA decay (NMD) is an mRNA surveillance pathway involved in translational control and gene expression regulation. Core NMD factors UPF1, UPF2 and UPF3B are conserved from yeast to humans and essential to target mcRNAs with a premature stop codon for decay. UPF2 binding to UPF1 activates UPF1’s ATPase and helicase activities, and UPF2 binding to UPF3B is important for its association with the exon-junction complex and efficient NMD. However, UPF2’s association with RNA remains largely uncharacterized. Here, we analyze nucleic acid binding, identifying the first and third MIF4G domains of UPF2 as main RNA-/DNA-binding modules. We find that UPF2’s MIF4G domain-3 has RNA annealing activity while full-length UPF2 unfolds our reporter hairpin-RNA structure. We show that UPF2 preferentially binds and stabilizes single-stranded RNA (ss-RNA) in a sequence-independent manner. Concomitant to ss-RNA binding, UPF2 undergoes a distinct conformational change in its otherwise highly dynamic structure. UPF2’s RNA binding and unfolding activity may support UPF1’s helicase and mRNP remodeling activity and, in combination with UPF3B, stabilize UPF1’s association with nonsense mRNA.

## INTRODUCTION

Nonsense-mediated mRNA decay (NMD) recognizes and degrades mRNAs with a premature termination codon (PTC) arising from genetic mutations, gene expression errors or alternative splicing events (Karousis and Mühlemann 2019; Kurosaki et al. 2019; Lejeune 2022; Tan et al. 2022). Thereby, NMD protects cells from the potentially harmful effects of C-terminally truncated protein products. Translation termination at a PTC is recognized by the NMD machinery which includes the key NMD factors Up-Frameshift Proteins UPF1, UPF2 and UPF3B. Assembly of the NMD machinery allows phosphorylation of UPF1 by the suppressors with morphological effects on genitalia protein 1 (SMG1) kinase complex. Phospho-UPF1 serves as a binding platform for the SMG5-SMG7 heterodimer and SMG6 endonuclease which cleaves the mRNA near the PTC, initiating decay. SMG5-SMG7 recruit mRNA decapping and deadenylation enzymes, as well as exonucleases, to degrade the mRNA. In addition to its role in mRNA quality control, NMD also controls 5-20% of human endogenous transcripts, thus regulating gene expression and shaping essential biological processes in development, differentiation and stress (Jaffrey and Wilkinson 2018; Nasif et al. 2018; Kurosaki et al. 2019). Loss or inactivation of NMD factors UPF1, UPF2, UPF3B, SMG1 and SMG6 cause lethality during early embryo development (Chousal et al. 2022). Importantly, NMD is implicated in ∼20% of human genetic diseases caused by a single base-pair mutation (Mort et al. 2008), and mutations in NMD factors are associated with neurodevelopmental disorders and various cancers (Jaffrey and Wilkinson 2018; Tan et al. 2022).

UPF1 is the key NMD factor involved in NMD substrate recognition, recycling of terminating ribosomes, remodeling of messenger-ribonucleoprotein complexes (mRNP) and initiation of decay (Fiorini et al. 2015; Lee et al. 2015; Serdar et al. 2020; Chapman et al. 2022). UPF1 binds RNA in a non-sequence-specific manner with high affinity when adopting a closed conformation (Cheng et al. 2007; Hogg and Goff 2010; Lee et al. 2015; Karousis and Mühlemann 2019). Binding of UPF2 to UPF1 stabilizes an open conformation of UPF1 with lower mRNA binding affinity, while enhancing UPF1’s RNA helicase and ATPase activities (Clerici et al. 2009; Chakrabarti et al. 2011). Thanks to its helicase activity, UPF1 translocates along the mRNA in a 5’ to 3’ direction, unwinds mRNA structures and dissociates or displaces RNA-bound proteins (Fiorini et al. 2015; Lavysh and Neu-Yilik 2020).

NMD factors UPF2 and UPF3B are associated with the exon-junction complex (EJC), which is deposited during mRNA splicing 20-24 nucleotides (nt) upstream of the exon-exon junction (Melero et al. 2012; Le Hir et al. 2016; Schlautmann and Gehring 2020). The presence of one or more EJCs in the 3’ untranslated region of an mRNA is a strong marker for NMD (Lindeboom et al. 2016). UPF2 and UPF3B are often presented as ‘adaptor proteins’, bridging UPF1, the SMG1-8-9 kinase complex and the EJC to form a decay-inducing (DECID) complex (Kashima et al. 2006). While both factors are essential for efficient NMD, UPF2-independent and UPF3B-independent NMD pathways have been reported (Gehring et al. 2005; Chan et al. 2009; Gehring et al. 2009; Mabin et al. 2018).

UPF2 is a 140 kDa perinuclear protein that comprises three middle portions of eIF4G (MIF4G) domains and a C-terminal UPF1-binding domain (U1BD) (Fig. 1A). The U1BD adopts a mixed α-helical/β-hairpin when interacting with UPF1’s cysteine-histidine-rich (CH) domain (Clerici et al. 2009). UPF2 U1BD binding induces an open conformation of UPF1 with enhanced ATPase and RNA helicase activities (Clerici et al. 2009; Chakrabarti et al. 2011). The third MIF4G domain (MIF4G-D3) of UPF2 is essential for NMD machinery assembly via a tight interaction with UPF3B which supports UPF2 association with the EJC (Kadlec et al. 2004; Buchwald et al. 2010; Bufton et al. 2022). MIF4G-D3 further binds to SMG1, facilitating the activation of SMG1 kinase activity (Kashima et al. 2006; Clerici et al. 2014; Deniaud et al. 2015). UPF2 MIF4G-D3 domain also binds RNA (Kadlec et al. 2004; Bufton et al. 2022), The UPF2 MIF4G-D3 domain also binds RNA (Kadlec et al. 2004; Bufton et al. 2022), whereas, to the best of our knowledge, DNA binding by UPF2 has not been reported. In fact, UPF2 binding to UPF3B induces a conformational change in UPF3B, interfering with RNA-induced oligomerization of UPF3B and delays translation termination by UPF3B (Neu-Yilik et al. 2017; Bufton et al. 2022). More recently, low-affinity RNA binding was shown for a construct comprising UPF2’s MIF4G domain-1 and domain-2 (MIF4G-D1-2) while no RNA binding was detected for the U1BD alone, indicating that the three MIF4G domains are responsible for RNA binding (Xue et al. 2023). Notably, this study also showed that RNA binding by UPF1 is weakened through interactions with UPF2, thereby preventing formation of a stable ternary complex (Xue et al. 2023).

**Figure 1.**
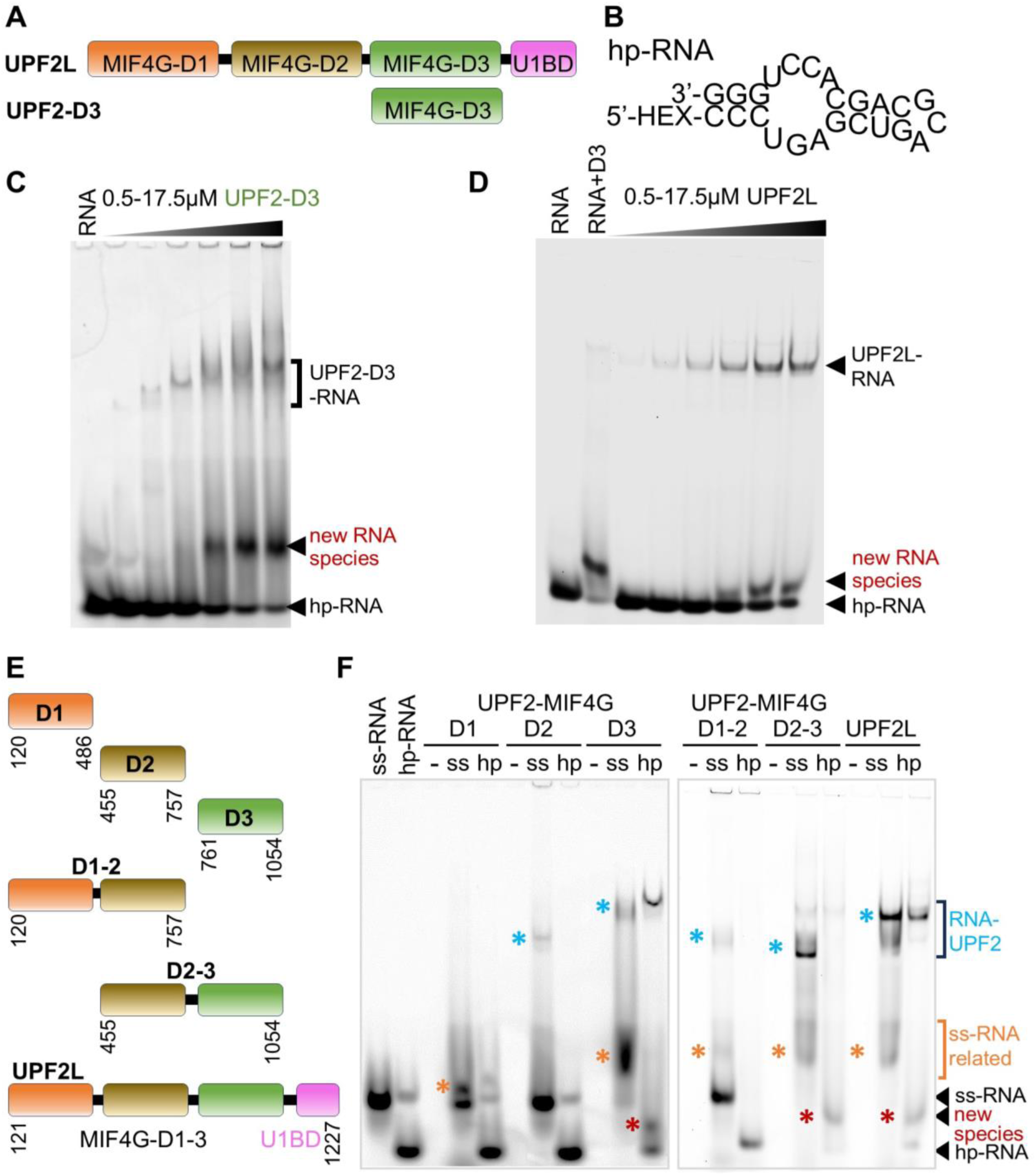
RNA binding of UPF2 variants tested by electromobility shift assays. **(A)** Schematic showing UPF2L (residues 121-1227) and UPF2 MIF4G-domain 3 (D3, 761-1054) constructs. **(B)** Hexachlorofluorescein (HEX)-labelled hairpin RNA (hp-RNA) construct used in this study (prediction based on RNAfold web server (Lorenz et al. 2011)). **(C-D)** EMSA using hp-RNA and increasing concentrations of MIF4G-D3 and UPF2L. New RNA bands appearing in the presence of UPF2 variants are highlighted (red). **(E)** UPF2 constructs used for EMSA shown in panel F. **(F)** EMSA using no RNA (-), ss-RNA (ss) and hp-RNA (hp) and UPF2 constructs. Corresponding Coomassie-stained gel sections are shown in Supplemental Fig. S1C. New RNA-containing bands are highlighted with asterisks.

Here, we systematically probed UPF2’s ability to bind nucleic acids. UPF2 and the MIF4G-D3 both bind RNA and DNA in their single-stranded (ss) and double-stranded (ds) forms. However, UPF2 has a clear preference for ss-RNA. Based on fluorescence anisotropy measurements, RNA binding by UPF2 is mostly mediated by MIF4G domains 1 and 3 (D1 and D3), with no clear preference for a particular sequence, pyrimidines or purines. Intriguingly, when analyzing UPF2’s binding to a model hairpin-RNA (hp-RNA) using electromobility shift assays (EMSA), we discovered that interaction between UPF2 or MIF4G-D3 with hp-RNA changes the RNA conformation. This activity is maintained in the presence of UPF1 and UPF3B. Notably, stable RNA associations are only formed by UPF2-UPF3B and UPF1-UPF2-UPF3B complexes. Using circular dichroism (CD) and fluorescence spectroscopy, we corroborate that UPF2 can unfold the hp-RNA. While UPF2’s structure is highly dynamic in the absence of RNA, binding of ss-RNA leads to more compact conformations of UPF2, reminiscent of the conformation reported previously after glutaraldehyde crosslinking (Melero et al. 2012; Lopez-Perrote et al. 2016). Our results suggest that the preferential binding to ss-RNA by UPF2’s MIF4G domains, in combination with UPF1-UFP2-UPF3B complex formation, enhances UPF1-UPF2’s attachment to nonsense RNA, contributes to RNA unfolding and mRNP remodeling, thereby facilitating mRNA decay.

## RESULTS

### UPF2 binds and remodels structured RNA

We previously reported RNA binding by the paralogues UPF3A and UPF3B using a 24-nt long 5’ hexachlorofluorescein (HEX)-labelled hp-RNA (Fig. 1B) (Neu-Yilik et al. 2017; Bufton et al. 2022). This RNA forms a hairpin with bulges, which likely mimics imperfect structures formed by mRNA more closely than a perfect hairpin or a poly(U) oligonucleotide which is single-stranded. A 24-nt RNA was used in this study as a longer RNAs show more conformational dynamics that enabled binding of two UPF2 molecules per RNA. Using EMSA, RNA-induced oligomerization of UPF3B was prevented by MIF4G domain-3 of UPF2 (Bufton et al. 2022). Instead, a stable complex comprising RNA, UPF3B and UPF2’s MIF4G-D3 was formed with defined stoichiometry (Bufton et al. 2022). Here, we tested the binding of UPF2 MIF4G-D3 alone with hp-RNA by EMSA. As expected, based on previous studies (Bufton et al. 2022; Xue et al. 2023), we observed the formation of hp-RNA/ MIF4G-D3 complexes (Fig. 1B,C). Surprisingly, addition of increasing concentrations of MIF4G-D3 led to the appearance of a defined, new RNA species giving rise to a band migrating slower in the gel than the hp-RNA input construct (Fig. 1C). Concomitantly, the hp-RNA band disappeared. This indicates that a different, distinct RNA structure is formed in the presence of high concentrations of UPF2’s MIF4G-D3.

Therefore, we next tested UPF2L (residues 121-1227), which comprises the three MIF4G domains and U1BD, and has the same activity as full-length UPF2 (Chakrabarti et al. 2011). We asked whether UPF2L would also induce the formation of a new RNA species in EMSA gels. A band corresponding to a defined UPF2L/ hp-RNA complex was observed with increasing concentrations of UPF2L (Fig. 1D). In addition, a new ‘RNA-only’ band appeared with increasing concentrations of UPF2L, with the hp-RNA band becoming less pronounced. This new RNA species migrated slower in the gel compared to hp-RNA alone (Fig. 1D). In fact, the position of the new RNA band was not identical to the band observed in the presence of MIF4G-D3, which ran higher in comparison (Fig. 1D). This suggests that a novel RNA structure is formed and released by UPF2L, which is different from the hp-RNA and the structure promoted in the presence of MIF4G-D3 alone. It also implies that the other domains of UPF2 contribute to RNA binding and rearranging the RNA structure.

### MIF4G domains mediate RNA binding of UPF2

We set out to determine which domains of UPF2 were responsible for RNA binding and forming these new RNA species. In addition to the 24-nt hp-RNA, we tested a 24-nt single-stranded ss-RNA construct derived from the hp-RNA by removing self-complementarity (Supplemental Fig. S1A). The design of the ss-RNA sequence was verified using RNA structure prediction software RNAfold (Lorenz et al. 2011), 3dRNA/DNA (Zhang et al. 2022) and trRosettaRNA (Wang et al. 2023) (Supplemental Fig. S1A-B). To identify the UPF2 domains responsible for RNA binding, we expressed six different UPF2 variants (Fig. 1E), comprising the individual MIF4G domains (D1, D2, D3), combinations of domains 1 and 2 (D1-2) and domains 2 and 3 (D2-3), as well as UPF2L. We did not test UPF2’s U1BD which does not bind RNA as shown previously (Xue et al. 2023).

In EMSA experiments, the ss-RNA runs higher in the gel compared to hp-RNA (Fig. 1F). All UPF2 constructs tested, with the exception of MIF4G-D1, bound ss-RNA as evidenced by protein-RNA complex bands for D2, D3, D1-2, D2-3, and UPF2L constructs (Fig. 1F, blue stars). Additionally, we observed RNA bands (discrete or diffuse) running higher in the gel for constructs D1, D3, D1-2, D2-3 and UPF2L (Fig. 1F, orange stars). This is consistent with complexes between ss-RNA and UPF2 variants dissociating during EMSA. A defined complex band was observed for hp-RNA/ UPF2L and hp-RNA/ MIF4G-D3, but not for the other UPF2 constructs (Fig. 1F). For the hp-RNA samples, we observed the formation of a new RNA species in the presence of MIF4G-D3, D2-3 and UPF2L, but not for the D1, D2 or D1-2 constructs (Fig. 1F, red stars). Therefore, it appears that RNA remodeling is specifically dependent on MIF4G domain-3 and only constructs comprising this domain (D3) could generate this novel RNA species.

Finally, ss-RNA or hp-RNA binding does not lead to a detectable upward shift of the protein bands in the EMSA gels for all UPF2 variants (Supplemental Fig. S1C). This indicates that no RNA-induced oligomerization or stable RNA/ UPF2 complexes occur under the conditions tested. Taken together, based on our EMSA experiments, it appears that all MIF4G domains of UPF2 contribute to RNA binding to varying degrees, with D3 being essential for RNA structural rearrangements.

### RNA complex formation and remodeling in the presence of UPF3B and UPF1

UPF1 and UPF3B are central protein interaction partners of UPF2. Therefore, we asked if the RNA binding and remodeling activity of UPF2 is maintained in the presence of these proteins. We previously established that a stable ternary complex comprising RNA, UPF3B and MIF4G-D3 can be formed (Bufton et al. 2022). To test for the presence of the new RNA species, we incubated UPF3B (residues 41-262 (Bufton et al. 2022)) with hp-RNA and added increasing amounts of MIF4G-D3 (Supplemental Fig. S2). In the absence of MIF4G-D3, a defined shift to the RNA-protein complex was observed upon addition of UPF3B, consistent with RNA binding (Hauer et al. 2016; Neu-Yilik et al. 2017; Bufton et al. 2022). However, adding a large excess of UPF3B compared to hp-RNA leads to RNA-induced oligomerization (Bufton et al. 2022). The RNA-UPF3B complex is too large to migrate into the gel and is found in the loading well (Supplemental Fig. S2). UPF3B’s oligomerization is prevented by adding increasing amounts of UPF2 MIF4G-D3, leading to the formation of defined MIF4G-D3/ UPF3B/ hp-RNA and MIF4G-D3/ hp-RNA complexes (Supplemental Fig. S2). In addition to these defined complexes, we observe a small hp-RNA band. The ‘new RNA’ species is detected in the presence of UPF3B and UPF2 MIF4G-D3 and UPF2 MIF4G-D3 alone, but not present in UPF3B alone (Supplemental Fig. S2). However, this new RNA species is not detected in the presence of UPF3B alone (Supplemental Fig. S2). Thus, UPF2 MIF4G-D3 appears to be active in the presence of UPF3B in remodeling the hp-RNA structure.

Next, we tested UPF2L’s RNA binding/ remodeling in the presence of UPF3B. Again, a new RNA species is formed with UPF2L and a UPF2L/UPF3B heterodimer (Fig. 2A). We detect the appearance of the ‘new RNA species’ with UPF2L alone and in the presence of UPF2L/UPF3B complex. As more RNA is bound by UPF2L/ UPF3B proteins, the hp-RNA band and the new RNA species become weaker but do not disappear entirely, despite excess of protein. We conclude that UPF2L remains active in RNA remodeling in the presence of UPF3B. Concomitantly, the affinity to RNA appears to be highest for UPF3B, and higher for the UPF2L/ UPF3B heterodimer compared to UPF2L alone, leading to a significant but not complete up-shift of the RNA in the EMSA gels.

**Figure 2.**
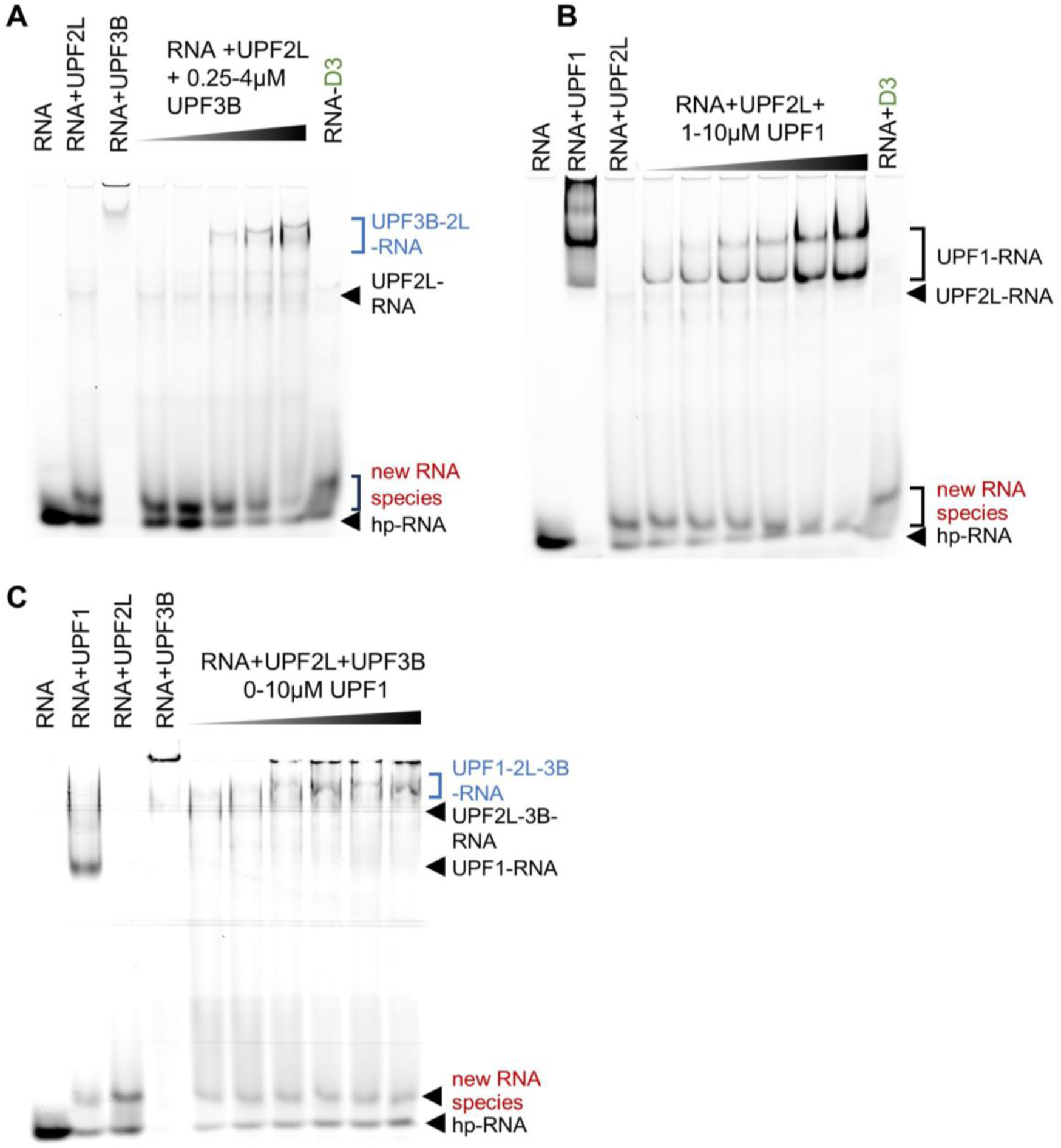
UPF2L complexes and RNA remodeling in the presence of UPF3B or UPF1. **(A)** EMSA using 250 nM hp-RNA in all lanes. As control, hp-RNA was incubated with 10 µM UPF2L alone or 4 µM UPF3B alone. A premix of 10 µM UPF2L with hp-RNA was supplemented with increasing concentrations of UPF3B. **(B)** EMSA using 250 nM hp-RNA in all lanes incubated with 10 µM UPF1 alone or 10 µM UPF2L alone. A premix of hp-RNA with 10 µM UPF2L was supplemented with increasing concentrations of UPF1. **(C)** EMSA using 250 nM hp-RNA in all lanes incubated with 10 µM UPF1 alone, 10 µM UPF2L alone, or 1.6 µM UPF3B alone. A premix of hp-RNA with 10 µM UPF2L and 4 µM UPF3B was supplemented with increasing concentrations of UPF1. **(A-C)** New RNA bands appearing in the presence of UPF2L or MIF4G domain-3 (control) are highlighted in red; RNA-protein complexes comprising UPF2L are highlighted in blue.

We then tested if the binding of UPF1 to UPF2L would interfere with UPF2’s RNA remodeling activity. UPF1 alone bound the RNA efficiently as evidenced by a complete up-shift of the hp-RNA band (Fig. 2B). UPF2L incubation with hp-RNA led to the appearance of the ‘new RNA species’ as observed previously (Fig. 1D). Addition of increasing amounts of UPF1 to UPF2L/ RNA resulted in an increased up-shifting of the RNA into an UPF1/ RNA complex. This affected both the hp-RNA band and the new RNA species band which decreased in intensity on the gel but did not disappear completely (Fig. 2B). A UPF2L/ UPF1/ RNA complex was not observed. Notably, UPF1 appears to have decreased affinity for RNA in the presence of UPF2L (Fig. 2B). These observations agree with previous reports showing that UPF2 destabilizes RNA binding by UPF1, confirming that ternary UPF2L/ UPF1/ RNA complexes are unstable and transient in nature (Chamieh et al. 2008; Xue et al. 2023).

Finally, we tested RNA in the presence of all three UPF proteins. UPF2L and UPF1 incubation with hp-RNA led to the appearance of the ‘new RNA species’ (Fig. 2C). Also, UPF3B-UPF2L-RNA complexes were formed, resolving the large RNA-UPF3B oligomers. Addition of increasing concentrations of UPF1 to UPF2L/ UPF3B-RNA complexes resulted in an increased up-shifting of the RNA into a UPF1/ UPF2L/ UPF3B-RNA complex. At the same time, the hp-RNA band and the ‘new RNA species’ band remained constant (Fig. 2C).

Taken together, the addition of UPF3B, or UPF3B and UPF1, to UPF2L leads to increased RNA-protein complex formation but free hp-RNA and the new RNA species can still be detected. This suggests that UPF2L can remodel hp-RNA in the presence of UPF1 and UPF3B, and that UPF3B is required for stable complexes containing RNA and UPF2.

### UPF2L preferentially binds single-stranded RNA

To quantify the contribution of the individual domains of UPF2 and understand the nature of UPF2’s interaction with nucleic acids, we analyzed DNA- and RNA-binding by fluorescence anisotropy. We first determined the affinity of the MIF4G domains alone and in combinations (Fig. 1E) for ss-RNA (Supplemental Fig. S1B). The dissociation constants (K_D_) for UPF2 MIF4G-D1, D2 and D3 constructs were determined as 1.4 µM, 3.5 µM and 694 nM, respectively (Fig. 3A). This indicates that D1 and D3 are mainly responsible for RNA binding with D3 being the key domain. UPF2 MIF4G-D1-2 and D2-3 bound ss-RNA with a K_D_ of 784 nM and 530 nM respectively. This is consistent with a minor contribution of D2 towards binding of ss-RNA by MIF4G-D3. In D1-D2, a more significant increase in affinity for binding ss-RNA is detected, indicating some cooperativity between D1 and D2 in RNA binding (Fig. 3A). Combining the binding sites of all three MIF4G domains, UPF2L had an affinity of 135 nM for ss-RNA. These K_D_ values are in good agreement with previously determined K_D_ values of 609 nM for a MIF4G-D3-U1BD variant and 761 nM for a D1-2 variant using a 12-nt poly(U) RNA (Xue et al. 2023). The agreement between the previously determined K_D_ values and our results also indicates that UPF2’s ss-RNA binding is sequence-independent. Taken together, these experiments show that in UPF2, all three MIF4G domains can interact with RNA, resulting in a K_D_ value of 135 nM which is compatible with mRNA binding by UPF2 in cells.

**Figure 3.**
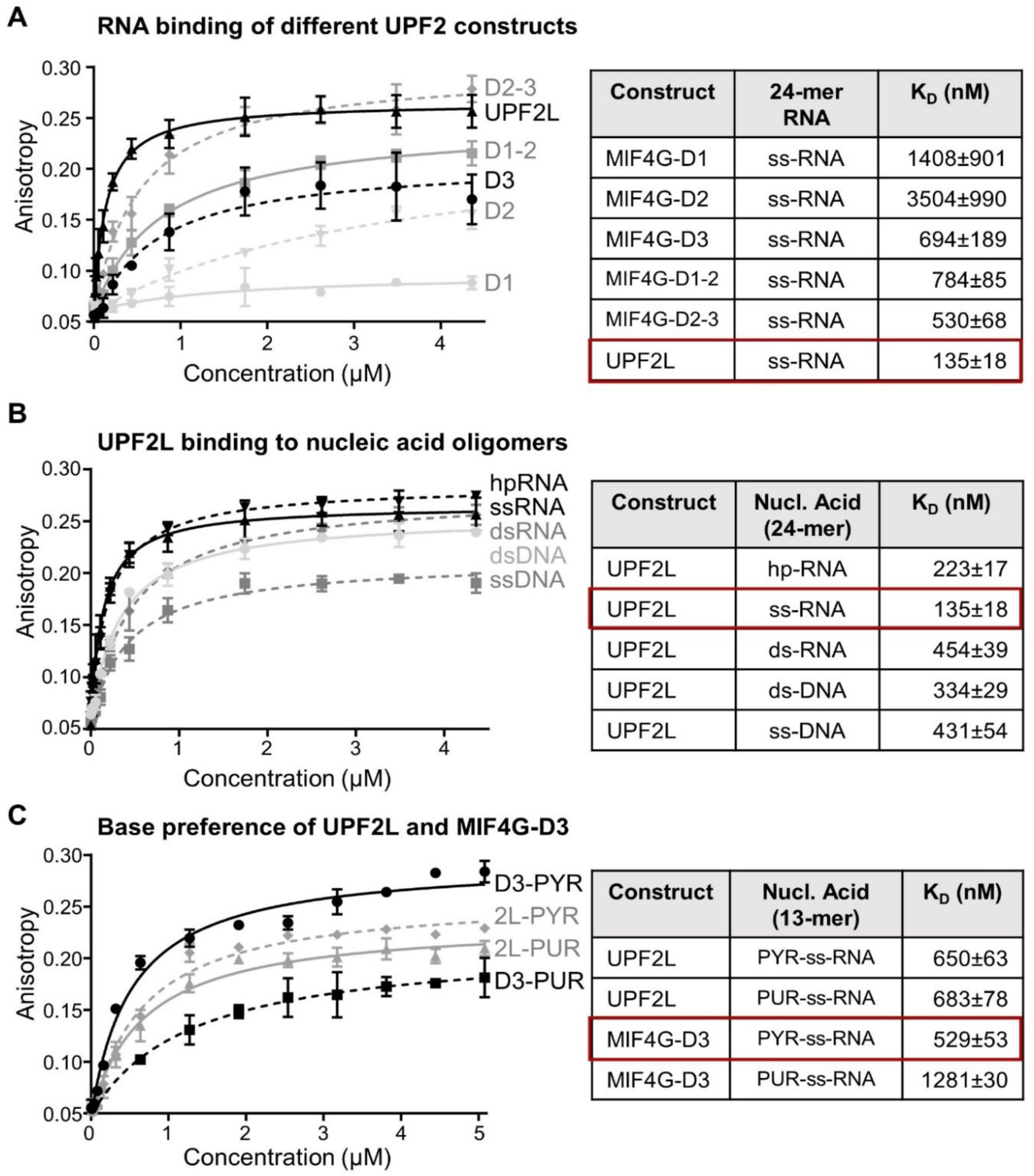
Nucleic acid binding of UPF2 variants determined by fluorescence anisotropy. **(A)** Fluorescence anisotropy binding curves of UPF2 constructs to HEX-labelled 24-nt ss-RNA (left). Measurements were performed in triplicate and error bars plotted via standard deviation before fitting a single component binding equation to calculate K_D_ values (right). **(B)** Binding curves of UPF2L to HEX-labelled 24-nt single-stranded (ss) and double-stranded (ds) RNA and DNA constructs (left) **(C)** Binding curves of UPF2L and MIF4G-D3 to HEX-labelled 13-nt pyrimidine-rich (PYR) or purine-rich (PUR) ss-RNA constructs (left). **(A-C)** K_D_ values are listed in the table (right) with highest affinity highlighted by a red box. Sequences are listed in Supplemental Table S1.

Subsequently, we investigated if UPF2 preferentially binds RNA or DNA in their double-stranded or single-stranded forms. The sequences of the nucleic acids used for fluorescence anisotropy are listed in Supplemental Table S1. First, we tested the 24-nt hp-RNA, the 24-nt ss-RNA and a 24-nt ds-RNA. Fluorescence anisotropy experiments reveal a clear difference in the binding affinities between ds-RNA and ss-RNA binding, with K_D_ values of 454 nM and 135 nM respectively (Fig. 3B). As expected, the hp-RNA, which comprises both double-stranded regions and single-stranded bulges (Supplemental Fig. S1A-B), bound with intermediate affinity to UPF2L (K_D_ of 223 nM, Fig. 3B). The difference in affinity between UPF2L binding to ds-DNA and ss-DNA (sequences in Supplemental Table S1) was less obvious, resulting in K_D_ measurements of 334 nM and 431 nM, respectively. This indicates a slight preference of UPF2 for binding ds-DNA (Fig. 3B). Clearly, UPF2L preferentially binds to ss-RNA as evidenced by its highest affinity (135 nM) for this construct in our experiments.

Next, we asked if we could identify sequence preferences for UPF2L-RNA binding. We tested two 13-nt RNA sequences that were either purine-rich or pyrimidine-rich (Supplemental Table S1). We could not identify a significant difference in RNA binding of UPF2L by fluorescence anisotropy as the pyrimidine-rich and purine-rich sequences bound RNA with very similar K_D_ values of 650 nM and 683 nM, respectively (Fig. 3C). The lower affinity of UPF2L towards these RNA oligonucleotides was due to the shorter RNA oligonucleotide tested; 13-nt versus 24-nt. In contrast, MIF4G-D3 displayed a strong preference for pyrimidine-rich RNA with a K_D_ of 529 nM, versus a K_D_ of 1.28 µM for purine-rich RNA (Fig. 3C). Here, the difference in K_D_ values for MIF4G-D3 ss-RNA binding between the two experiments (694 nM versus 529 nM for the 13-nt versus 24-nt ss-RNA) appears to originate from the different sequences and not the length of the oligonucleotide. A significant decrease in affinity for shorter RNAs (<13 nt) was observed for MIF4G-D3, with very weak binding to a 9-mer RNA oligonucleotide (Supplemental Fig. S3). This suggests that high-affinity RNA binding by UPF2 MIF4G-D3 requires an RNA sequence length of at least 13-nt with a preference for pyrimidine-rich sequences.

In summary, UPF2 can bind DNA and RNA in their single-stranded and double-stranded form. UPF2 preferentially binds ssRNA oligonucleotides longer than 13 nucleotides, and we could not identify any sequence preference or preference for purines or pyrimidines for UPF2’s nucleic acid binding.

### UPF2 remodels structured RNA

In our EMSA experiments we observed the appearance of a new band of RNA in the presence of UPF2L and MIF4G-D3 (Fig. 1C,D). However, it remained unclear whether the new band corresponds to single-stranded, double-stranded or alternatively/ partially folded RNA. Unfolding of the RNA hairpin would result in ss-RNA which is expected to migrate higher in a gel due to its less compact conformation. Given the high complementarity between hp-RNA sequences, UPF2L could also anneal two hp-RNA molecules to form ds-RNA (Fig. 4A). In EMSA, ds-RNA is also expected to run higher in the gel compared to intramolecularly folded hp-RNA. Annealing of complementary RNAs can occur spontaneously or be accelerated by a protein (Fig. 4A). Such a ‘protein annealer’ can bind one or both RNAs, alter the RNA structure, enhance the local concentration of RNA and thus the probability of ds-RNA formation (Rajkowitsch et al. 2007). In fact, the three MIF4G domains of UPF2 offer at least two RNA-binding sites in MIF4G-D3 and D1-D2 (Fig. 3A). By binding two RNA molecules, likely in their single-stranded form, UPF2 could thus facilitate annealing of two RNA molecules into a single ds-RNA due to their proximity.

**Figure 4.**
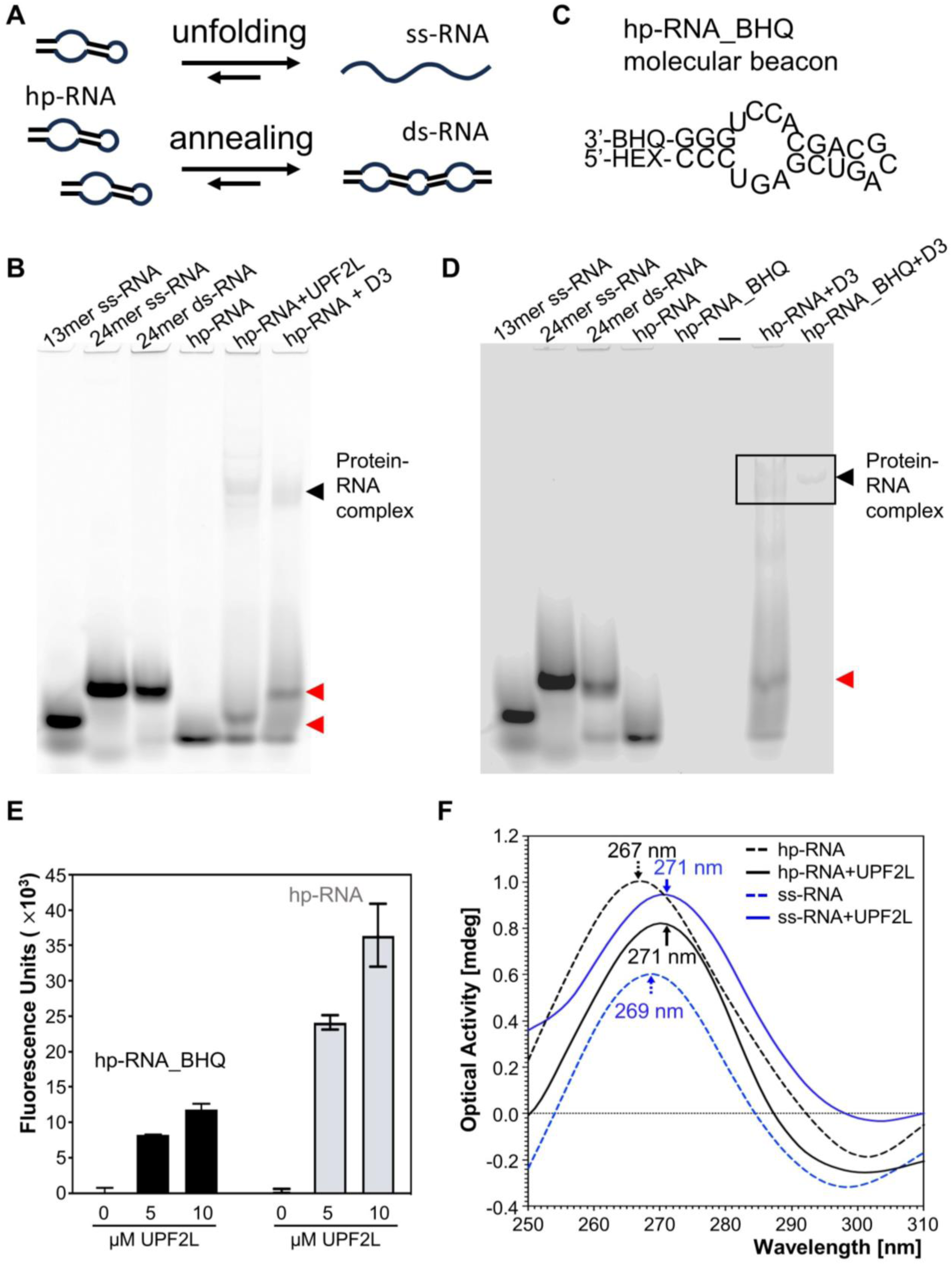
RNA structural changes mediated by UPF2L. **(A)** A schematic illustrating potential UPF2-mediated RNA annealing or RNA unfolding activities. **(B)** EMSA using ss-RNA oligonucleotides of 13 nucleotides, 24 nucleotides, ds-RNA and hp-RNA as controls. Incubation of hp-RNA with UPF2L or MIF4G-D3 produces new RNA bands running at similar height as 13-nt ss-RNA and 24-nt ss-RNA/ ds-RNA, respectively (red arrows). **(C)** Molecular beacon hp-RNA (hp-RNA_BHQ1) construct labelled with hexachlorofluorescein (HEX) dye and blackhole quencher 1 (BHQ1). **(D)** EMSA using same controls as in panel B, 250 nM molecular beacon, and MIF4G-D3 incubated with hp-RNA or molecular beacon. The new RNA band is highlighted (red arrow). **(E)** UPF2L-mediated RNA structural changes monitored by fluorescence spectroscopy using the molecular beacon (black bars) and hp-RNA (grey bars). Experiments were performed in duplicate. **(F)** Changes in hp-RNA and ss-RNA tertiary structure upon addition of 10 μM UPF2L monitored by CD spectroscopy (hp-RNA, black dashed line; ss-RNA, blue dashed line; hp-RNA spectrum subtracted by UPF2L spectrum, black line; ss-RNA spectrum subtracted by UPF2L spectrum, blue line). Peak wavelengths are indicated. For reference: single-stranded poly-uridine peaks at 272 nm.

To investigate this, we first performed EMSAs to test for unfolding or annealing. Using varying RNA lengths as size markers in EMSA gels, we loaded HEX-labelled 13-nt ss-RNA, 24-nt ss-RNA, 24-nt ds-RNA and the 24-nt hp-RNA (Fig. 4B). Incubation of the hp-RNA with UPF2L leads to a band that ran similar to the 13-nt ss-RNA in EMSA, whereas incubation with MIF4G-D3 resulted in a band running at approximately the same height of ss-RNA and ds-RNA (Fig. 4C). This suggests that MIF4G-D3 fully unfolds or anneals the hp-RNA. In contrast, UPF2L binding to hp-RNA results in a partially unfolded RNA species.

To discriminate between RNA annealing and RNA unfolding by UPF2L and MIF4G-D3, we tested a molecular beacon hp-RNA construct with a HEX dye at the 5’-end and a blackhole quencher 1 (BHQ-1) at the 3’-end (Fig. 4C) (Tyagi and Kramer 1996). This molecular beacon was used in EMSA and for fluorescence spectroscopy to determine changes in fluorescence upon addition of UPF2L. Unfolding of RNA is expected to increase the fluorescence of the molecular beacon as the HEX dye would no longer in close proximity to the BHQ-1 quencher. In contrast, RNA annealing of the hp-RNA should not alter the fluorescence signal, since the 3’ BHQ-1 of the annealed complementary RNA strand will quench the 5’ HEX dye.

We performed EMSA using the same size markers as before (Fig. 4B,D). As expected, we could not detect a signal for the molecular beacon, confirming quenching of the HEX dye. In the presence of MIF4G-D3, a weak band was observed for the molecular beacon at the height of the RNA-protein complex (Fig. 4D). This indicates that the molecular beacon hp-RNA is unquenched when bound to MIF4G-D3. No band was observed at the height of the new RNA species (Fig. 4D). This suggests that the molecular beacon is still quenched and thus in a double-stranded rather than single-stranded conformation. We conclude that MIF4G-D3 exhibits RNA annealing activity. The same experiment using UPF2L did not show any fluorescent signal for the UPF2L-RNA complex or RNA alone in the EMSA gel and thus was not conclusive (not shown). Therefore, we decided to determine any conformational change by using fluorescence spectroscopy which is an equilibrium measurement method and potentially more sensitive in detecting unquenched RNA.

Fluorescence spectroscopy would reveal any changes in the environment of the HEX dye in the molecular beacon and the hp-RNA constructs in the presence of UPF2L, leading to observed structural changes of the RNA in EMSA gels. In the absence of UPF2L, background fluorescence was detected, indicating that not all hp-RNA and molecular beacon probes are in a hairpin conformation (Supplemental Fig. S4). In agreement, we occasionally observed a second band for the ‘hp-RNA only’ control, running slower in the EMSA gel (Fig. 1C,F). Addition of increasing concentrations of UPF2L led to increased fluorescence signals suggesting spatial separation of the fluorescent dye and the BHQ-1 quencher in the molecular beacon, consistent with unfolding (Fig. 4E, Supplemental Fig. S4). Notably, we observed an increase in fluorescence of the hp-RNA upon addition of UPF2L (Fig. 4E). This is likely due to protein-induced fluorescence enhancement (PIFE), a photo-physical effect where binding of a protein in proximity leads to increased fluorophore intensity (Hwang and Myong 2014). We conclude that the observed fluorescence increase in the presence of UPF2L is due to a conformational change of the molecular beacon RNA structure to ss-RNA, and concomitant unquenching of the HEX dye, and PIFE.

To further confirm the formation of ss-RNA rather than ds-RNA in the presence of UPF2L, CD spectroscopy was performed, allowing detection of changes in tertiary RNA structure. In its A-form, ds-RNA is known to produce in CD spectra a positive peak at ∼260 nm and a negative peak near 210 nm, whereas ss-RNA (such as poly(U)) displays a positive peak at ∼272 nm (Chauca-Diaz et al. 2015; Szpotkowski et al. 2023). Our 24-nt ss-RNA displayed a positive peak at 269 nm, indicating it is not completely unstructured, conflicting with computational predictions (Fig. 4F, Supplemental Fig. S1A-B). The 24-nt hp-RNA containing double-stranded RNA and single-stranded bulges showed a peak at 267 nm in CD spectra (Fig. 4F). This peak value agrees with a small fraction of unfolded RNA. In the presence of excess UPF2L, we observed a shift of the ss-RNA peak from 269 nm to 271 nm after subtracting the CD curve of UPF2L protein only (Fig. 4F), in agreement with the ss-RNA peak value described in the literature. This indicates that UPF2L binding stabilizes the ss-RNA conformation. Similarly, a shift is observed for the hp-RNA from 267 nm to 271 nm in the presence of excess UPF2L (Fig. 4F) indicating formation of ss-RNA. In summary, the results from CD spectroscopy and molecular beacon fluorescence spectroscopy confirm that UPF2L opens hp-RNA structures and stabilizes single-stranded RNA.

### UPF2L structural changes upon RNA binding

We subtracted the CD curves of hp-RNA and ss-RNA from the CD curves of UPF2L/ hp-RNA and UPF2L/ ss-RNA, respectively, to identify any changes in UPF2L structure upon RNA binding. Notably, the CD spectra for UPF2L alone, UPF2L + hp-RNA and UPF2L + ss-RNA showed clear differences between the three samples in the region of 210-230 nm (Supplemental Fig. S5). While all three spectra displayed a negative peak at a wavelength of ∼222 nm characteristic of α-helical secondary structure, the ellipticity intensity at the 222 nm peak was different for the three samples. Unexpectedly, a ∼3-times stronger negative signal was observed for the UPF2L + ss-RNA complex compared to UPF2L alone (Supplemental Fig. S5). The UPF2L + hp-RNA sample signal amplitude at 222 nm was approximately half of UPF2L alone (Supplemental Fig. S5). The high concentration of UPF2L required for the experiment (10 µM) precluded further characterization of the secondary structure change. Nonetheless, these results indicate that UPF2L changes its structure when binding RNA and becomes more α-helical when in a complex with ss-RNA.

### Single-stranded RNA binding induces compact UPF2L conformations

To further investigate the structural changes in UPF2L upon RNA binding observed by CD spectroscopy, we performed negative-stain electron microscopy (EM) and 2D classification (Fig. 5A-C, Supplemental Fig. S6A-C). In the 2D class averages of UPF2L alone we observed a huge variability in shapes, ranging from elongated L and S shapes to more compact V and U shapes, including a closed (doughnut-like) shape (Fig. 5A). The dimensions of the particles range from 10 nm for the more compact shapes to almost 20 nm for the elongated shapes. A similar variation in sizes and shapes was observed for the UPF2L sample which was incubated with a 4-fold molar excess of hp-RNA (Fig. 5B), indicating a high degree of flexibility of the protein. In contrast, incubation of UPF2L with 4-fold molar excess of ss-RNA led to more defined particles in the micrographs (Supplemental Fig. S6C) and less variability in the 2D class averages (Fig. 5C). Most particles adopt U- or V-shaped conformations in the presence of ss-RNA. In further support of this observation, we quantified the number of particles found in 2D classes showing L/S-shaped, elongated particles versus U/V-shaped, compact particles (Fig. 5A-C, Supplemental Fig. S6D). For UPF2L alone and UPF2L/ hp-RNA samples, ∼90% of the particles were found to be elongated. In contrast, in the UPF2L sample incubated with ss-RNA, ∼92% of particles adopted compact conformations (Fig. 5A-C, Supplemental Fig. S6D). Remaining flexibility of the more compact UPF2L/ ss-RNA particles precluded a high-resolution cryo-EM study. The compact U/V-shaped particles observed in the current work resemble previously reported low-resolution UPF2 cryo-EM structures where UPF2 was crosslinked with glutaraldehyde prior to single-particle analysis (Melero et al. 2012; Lopez-Perrote et al. 2016). Taken together, our negative-stain EM results and our observations from CD spectroscopy both show that UPF2L changes its structure upon binding ss-RNA. More compact conformations are adopted in complex with ss-RNA, reminiscent of glutaraldehyde-crosslinked UPF2.

**Figure 5.**
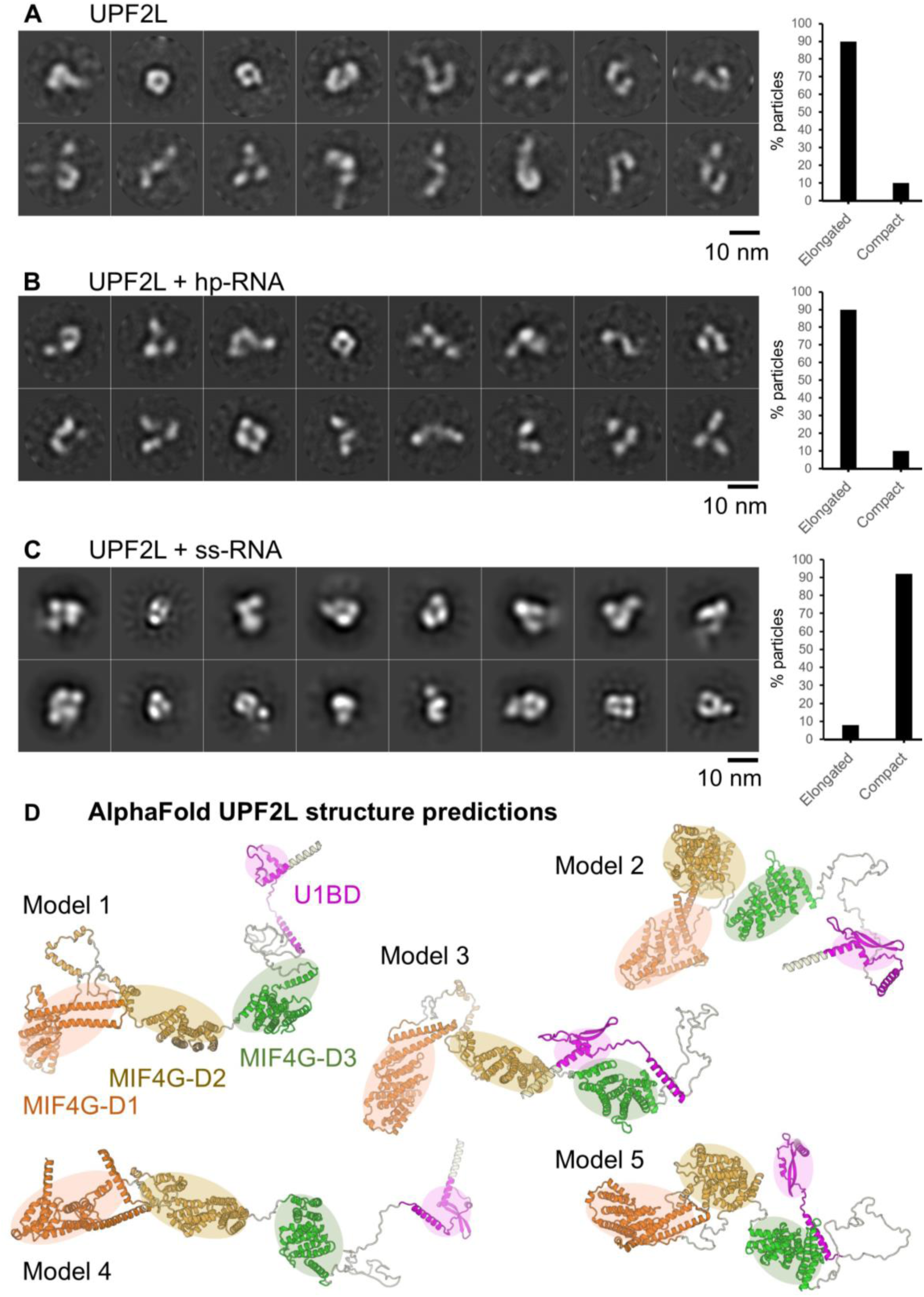
Dynamics and conformational change of UPF2L upon RNA binding. **(A)** Negative-stain EM 2D class averages of UPF2L alone. **(B)** Negative-stain EM 2D class averages of UPF2L with hp-RNA. **(C)** Negative-stain EM 2D class averages of UPF2L with ss-RNA, leading to more defined, compact particles. **(A-C)** Scale bar: 10 nm. Right: Plot showing the percentage of particles classifying into elongated versus compact, U/V-shaped 2D classes. **(D)** Relaxed AlphaFold2 models of UPF2L (without RNA) showing highly dynamic regions between the individual MIF4G domains and the UPF1-binding domain (D1: orange, D2: brown, D3: green, U1BD: magenta), agreeing with conformational dynamics and mostly elongated particles observed in panels A and B.

We note that several UPF2L dimers with compact conformations were also observed in the sample with ss-RNA in negative-stain EM and 2D class averages, despite an excess of ss-RNA (Supplemental Fig. S7A). We wondered if the 24-nt ss-RNA could accommodate the binding of multiple UPF2L molecules. However, our K_D_ measurements indicate that shorter RNAs are bound by UPF2L with lower affinity (13-nt, Fig. 3C). Therefore, we asked whether two ss-RNA molecules could interact by partial annealing. For the computational analysis, we artificially linked two molecules via a U_10_ or A_10_ linker. RNAfold (Lorenz et al. 2011), MXfold2 (Sato et al. 2021) and 3dRNA/DNA (Zhang et al. 2022) predict the formation of a short RNA-helix and thus possible dimerization, whereas trRosettaRNA (Wang et al. 2023) does not predict any interaction between the ss-RNA molecules (Supplemental Fig. S7B). We conclude that the UPF2L/ ss-RNA dimers observed in EM are likely mediated by RNA binding and by UPF2L alone.

Finally, AlphaFold2 (Mirdita et al. 2022) and Rosetta Relax (Tyka et al. 2011) were used to predict the structure of the UPF2L protein (Fig. 5D). Predicted models highlight a flexible, rather elongated arrangement of the MIF4G domains and U1BD (Fig. 5D), recapitulating the L- and S-shapes of UPF2L observed in the 2D class averages from negative-stain EM (Fig. 5A). Taken together, our results indicate a flexible domain arrangement of UPF2L both in the absence of RNA and in the presence of structured RNA (hp-RNA), which shifts to more compact conformations when a complex is formed with ss-RNA.

## DISCUSSION

In this study, we analyzed RNA binding by UPF2 and its domains. Our experiments show that UPF2L can bind DNA but preferentially binds single-stranded RNA with no apparent sequence specificity. We discovered that UPF2L and MIF4G-D3 alter the structure of hairpin RNA. Notably, the novel RNA species observed in EMSA are not identical for these proteins. The new RNA species produced by MIF4G-D3 migrates slower in the gel compared to the species observed in the presence of UPF2L. This indicates that additional domains of UPF2 also play a role in nucleic acid binding and remodeling. In fact, fluorescence anisotropy experiments demonstrate that all three MIF4G domains can interact with ss-RNA. MIF4G-D1 has an affinity of 1.4 µM, and MIF4G-D2 has a much weaker affinity of 3.5 µM. However, in combination, D1-D2 bind ss-RNA with an affinity of ∼800 nM. This is in agreement with the K_D_ for MIF4G-D1-2 determined in a previous study testing shorter U_15_ RNA (Xue et al. 2023). MIF4G-D3 alone binds the model RNA with ∼700 nM affinity, and in combination with D2 the K_D_ moderately increased to ∼530 nM, confirming a small contribution of D2. UPF2L binds ss-RNA with a K_D_ of 135 nM, in agreement with the avidity expected for at least three RNA-binding sites. This indicates that UPF2 is an RNA binding protein under physiological conditions, unless its binding to nucleic acids would be prevented by complex formation with other proteins.

When we tested UPF2’s RNA-interactions in the presence of UPF3B and UPF1 using EMSA, we found that a tertiary complex can be formed by UPF2L/ UPF3B/ hp-RNA and that an additional RNA species running higher than hp-RNA is still observed in the presence of UPF2L and UPF3B. Addition of UPF1 to UPF2L/ hp-RNA complexes led to increased UPF1/ RNA complex formation, However, the new RNA species is still observed, indicating that UPF2L can remodel the unbound hp-RNA. Notably in EMSA, the gel shift observed for the UPF1/ hp-RNA is significantly stronger than the shift observed for UPF1 in the presence of UPF2L and hp-RNA indicating decreased affinity for RNA. There is also no sign of a UPF1/ UPF2L/ hp-RNA complex in EMSA gels, confirming that UPF1 binds either UPF2 or RNA and that the ternary complex is unstable (Chamieh et al. 2008; Chakrabarti et al. 2011; Xue et al. 2023). When UPF1 was added to UPF2L/ UPF3B/ RNA complexes an additional band corresponding to a UPF1/ UPF2L/ UPF3B/ RNA complex was detected, indicating that UPF3B is important for stable RNA association (Fig. 2).

UPF2L does not form stable RNA-protein complexes in EMSA, a non-equilibrium method. Free single-stranded and hairpin RNAs are mostly released from UPF2L in EMSA, even in the presence of excess UPF2L. This also applies to all UPF2 variants tested. We suggest that UPF2L and MIF4G-D3 dynamically bind and release RNA. Concomitantly, they remodel the hp-RNA and release a new RNA species in a process that does not consume energy. Notably, the new RNA band observed in the presence of UPF2L runs at the same height as 13-nt ss-RNA and not at the height of 24-nt ss-RNA, suggesting that the RNA is not completely unstructured/ unfolded. In marked contrast, in the presence of MIF4G-D3 a band corresponding to 24-nt ss-RNA or 24-nt ds-RNA is detected by EMSA. This new RNA band is completely quenched when a molecular beacon probe is used, indicating that it corresponds to ds-RNA, while a small fraction of unquenched ss-RNA is detected in a MIF4G-D3/ RNA complex. We conclude that MIF4G-D3 exhibits an annealing activity, bringing two unstructured RNA molecules in close proximity to form ds-RNA. Dequenching of the molecular beacon was also observed in the presence of UPF2L in fluorescence spectroscopy (Fig. 4). This indicates that the HEX fluorescence label has a larger distance to the BHQ-1 quencher and, accordingly, the hp-RNA is unstructured in that region. This finding was further corroborated by CD spectroscopy showing an increased proportion of ss-RNA in the presence of UPF2.

CD spectroscopy indicated that UPF2L is highly flexible, even in the presence of hp-RNA. In contrast, an increase in secondary structure was observed in the presence of the 24-nt ss-RNA, suggesting UPF2L/ ss-RNA complex formation and a conformational change in the protein. Consistently, we observe that UPF2L and UPF2L/ hp-RNA adopt many different conformations in negative-stain EM 2D class averages (Fig. 5). Similarly, AlphaFold2 (Mirdita et al. 2022) and Rosetta Relax (Tyka et al. 2011) predict a very flexible domain arrangement with virtually no interactions between the individual domains of UPF2. In contrast, more compact U-/V-shaped UPF2L conformations are observed in the presence of ss-RNA in negative-stain EM 2D class averages. We note that the pseudo-atomic model of MIF4G domains 2 and 3 from crystallography (Clerici et al. 2014) could not be fitted into the negative-stain EM density of glutaraldehyde-crosslinked UPF2 (Melero et al. 2012), indicating different conformations. This further supports that UPF2 domains do show no interactions with each other and UPF2 is a highly dynamic protein.

We find that UPF2 preferentially binds and stabilizes unstructured RNA. Moreover, UPF2 can unfold weakly structured RNA, also in the presence of UPF3B. Given that UPF2 is a conserved NMD factor which interacts with and activates UPF1 helicase (Kadlec et al. 2004; Chakrabarti et al. 2011; Fiorini et al. 2015; Serdar et al. 2016; Serdar et al. 2020), this UPF2 activity is likely to be relevant in the context of mRNP remodeling and ultimately mRNA decay. It is conceivable that UPF2 binds unstructured mRNA after UPF1 helicase has unwound it (Fig. 6). Thus, UPF2 could prevent re-folding of mRNA by binding it and/ or contribute to unfolding mRNA structures, in synergy with UPF1 RNA helicase.

**Figure 6.**
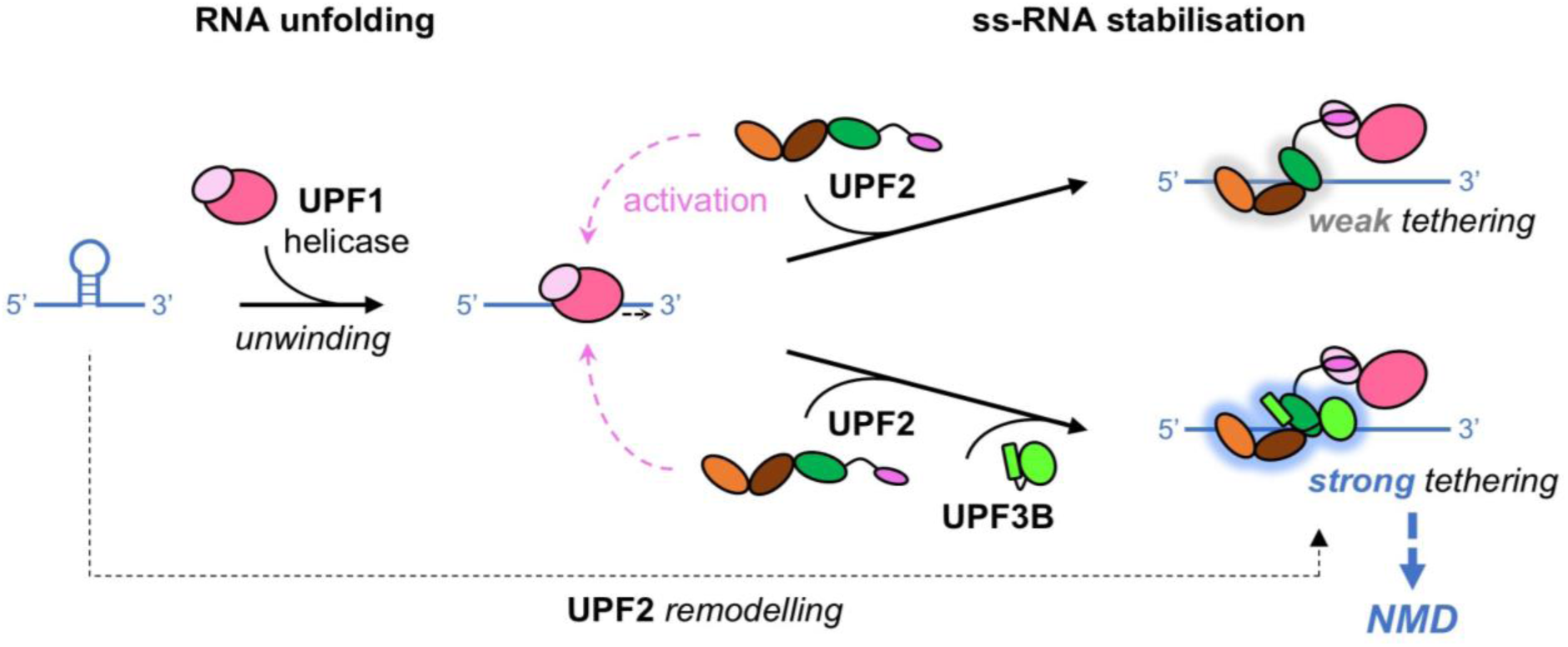
Model of UPF2-RNA interactions. Our data suggest a highly dynamic UPF2 structure and different RNA complex formation in the presence and the absence of UPF1 and/or UPF3B. UPF1 is activated by UPF2, and then weakly tethered to RNA. UPF3B can stabilize UPF1-UPF2‘s association with RNA, and UPF2 can support UPF1’s RNA unfolding activity, corroborating mRNA degradation.

Moreover, UPF2 and UPF3B provide additional RNA-binding sites to the NMD machinery, facilitating the association of UPF1 with a nonsense mRNA substrate. UPF2 also supports UPF1’s activation and phosphorylation by SMG1 kinase (Kashima et al. 2006; Clerici et al. 2009; Chakrabarti et al. 2011; Clerici et al. 2014; Deniaud et al. 2015). The additional RNA-binding sites of UPF2 are likely not sufficient to ensure that activated UPF1 remains associated with the nonsense RNA because the ternary complex is unstable (Fig. 2B, (Chamieh et al. 2008; Chakrabarti et al. 2011)). However, the combination of UPF2 and UPF3B leads to more stable ternary complexes with RNA (Fig. 2). UPF2/ UPF3B complexes could thus ensure a robust activation of UPF1 and its stable association with a nonsense mRNA (Fig. 6). In fact, the binding and ATPase-dependent dissociation of UPF1 from RNA was found to be important for NMD target discrimination (Lee et al. 2015; Chapman et al. 2022; Fritz et al. 2022). UPF2/ UPF1 complex formation and concomitant activation of UPF1 ATPase activity are suggested to contribute to protecting non-target mRNAs because the UPF2-induced destabilization of RNA binding could promote UPF1 dissociation and support efficient recycling of UPF1 from the non-target RNA (Chapman et al. 2024). Conversely, NMD factors should tether activated UPF1 to the target nonsense mRNA for efficient decay; reminiscent of tethering assays of NMD factors to the 3’UTR of reporter mRNAs which efficiently initiate NMD (Lykke-Andersen et al. 2000; Gehring et al. 2008). A stable UPF1 association after ATPase activation could be achieved by UPF2/ UPF3B complexes (Fig. 6). In fact, the MIF4G domains of UPF2 can bind RNA together with the N-terminal UPF3B domains (Fig. 2 and (Bufton et al. 2022)). We note that individual nucleotide resolution UV crosslinking and immunoprecipitation (iCLIP) experiments indicated that UPF3B can interact with RNA in complex with EJCs and in a non-EJC context (Hauer et al. 2016), the latter may be essential for EJC-independent NMD (Bühler et al. 2006; Metze et al. 2013). In summary, we propose that stable association of activated UPF1 with the nonsense RNA could be achieved by RNA binding of the UPF2/ UPF3B complex, alone or when associated to EJCs, to ensure efficient NMD; and UFP2 functions to support UPF1’s RNA helicase and mRNP remodeling activities.

## MATERIALS AND METHODS

### Protein production

pProExHTb plasmids (Invitrogen) encoding near full-length UPF2L (121-1227) (Neu-Yilik et al. 2017), MIF4G-D3 (761-1054) (Kadlec et al. 2004), MIF4G-D1 (120-486), MIF4G-D2 (455-757), MIF4G-D1-2 (120-757) and MIF4G-D2-3 (455-1054) constructs (Clerici et al. 2014) all comprise a C-terminal TEV-cleavage site and a His_6_-tag. The plasmids were transformed into *Escherichia coli* BL21 Rosetta (DE3) (Novagen) cells and grown in LB media supplemented with 100 μg/ml ampicillin to an OD_600nm_ of 0.6-0.8, induced with 1 mM IPTG (isopropyl 1-thio-β-D galactopyranoside) and incubated overnight (∼16 hours) at 20°C. After harvesting by centrifugation (5,000 x *g* at 4°C for 15 min), cell pellets were resuspended in 25 mM HEPES pH 7.45, 300 mM NaCl, 10 mM imidazole, 0.05% TWEEN 20, 5% glycerol supplemented with cOmplete EDTA-free protease inhibitor cocktail tablet (Roche) and Benzonase Nuclease (Merck). Cells were lysed by sonication and the cell lysate clarified by centrifugation at 45,000 x *g* at 4°C for 60 min. The supernatant was incubated with 5 ml Ni-NTA resin at 4°C for 1 hour. Beads were washed with 25 mM HEPES pH 7.45, 300 mM NaCl, 30 mM imidazole, 5% glycerol buffer. The proteins were eluted by incubation with 25 mM HEPES pH 7.45, 300 mM NaCl, 5% glycerol buffer containing increasing concentrations of imidazole from 50-250 mM. Eluted fractions containing the protein of interest were pooled and buffer-exchanged to 25 mM HEPES pH 7.45, 150 mM NaCl, 1 mM TCEP and 5% glycerol. The sample was applied onto a HiTrap Heparin column (Cytiva) using a 50 ml superloop (GE Healthcare). After washing, the proteins were eluted via a linear gradient of 150-1000 mM NaCl over 20 column volumes. Elution fractions were analyzed by SDS-PAGE. Fractions containing the protein of interest were pooled and concentrated to ∼8-10 mg/ml using an appropriate molecular weight cut-off concentrator (Merck). Concentrated protein fractions were further purified by size exclusion chromatography (SEC) using a Superdex 75 Increase 10/300 GL column or a Superdex S200 10/300 GL column (GE Healthcare), depending on the molecular weight of the UPF2 variant. Columns were equilibrated with SEC buffer (25 mM HEPES pH 7.45, 300 nM NaCl, 1 mM TCEP, 5% glycerol). Peak fractions were analyzed by SDS-PAGE, pooled and quantified using a Nanodrop One spectrophotometer (Thermo Scientific), followed by concentrating to ∼8-10 mg/ml. Extinction coefficient and molecular weight were determined using ExPASy: SIB bioinformatics resource portal (www.expasy.org). Concentrated protein was flash frozen in aliquots at -80°C for storage.

Full-length UPF1 was expressed using the MultiBac baculovirus-insect cell expression (Gupta et al. 2019) and purified as previously described (Neu-Yilik et al. 2017). UPF3B comprising residues 41-262, consisting of the RNA-recognition motif, NOPS-Like domain and the first coiled coil-like domain, was expressed in *E. coli* and purified as described (Bufton et al. 2022).

### Nucleic acid oligonucleotides

The sequences of the oligonucleotide constructs used in this study are listed in Supplemental Table S1. The DNA and RNA oligonucleotides are based on a 24-nt oligonucleotide sequence previously utilized (Neu-Yilik et al. 2017). HEX-labelled probes were ordered from Eurofins Genomics UK and dissolved using 100 μL of Milli-Q water to generate a high stock concentration. Oligomer concentrations were determined using a Nanodrop One spectrophotometer (Thermo Scientific). Double-stranded 24-nt DNA and RNA (ds-DNA / ds-RNA) was prepared by mixing the complementary oligonucleotides at a 1:1 molar ratio, heating to 65 ⁰C and slow cooling to room temperature overnight in a polystyrene container. Resulting stock concentrations of oligomers were aliquoted into small volumes and flash frozen at -80°C for storage.

### Electrophoretic mobility shift assay (EMSA)

EMSA experiments were performed using HEX-labelled RNA oligonucleotides (Supplemental Table S1) diluted to 250 nM in 25 mM HEPES pH 7.45, 100 mM NaCl, 5% glycerol. Excess concentration of probe was necessary to detect multiple band shift species (Bufton et al. 2022). Serial dilutions of UPF2L, MIF4G-D3, UPF3B (residues 41-262) and/or UPF1 was performed using buffer containing 25 mM HEPES pH 7.45, 300 nM NaCl, 5% glycerol, and 2 mM β-mercaptoethanol or 1 mM TCEP. Proteins, RNA and native sample buffer (Thermo Scientific) were mixed and incubated on ice for 1 hour before loading onto Native Novex WedgeWell 4-20% Tris-Glycine gels in Tris-glycine native running buffer (Invitrogen). Gels were run for 45 min at 150 V at 4°C before detecting the HEX-label using a Typhoon FLA 9500 instrument (GE Healthcare) at 532 nm wavelength and BPG1 emission filter. EMSA gels were stained for protein using Coomassie Brilliant Blue.

### Circular dichroism (CD) spectroscopy

The 24-nt oligonucleotides ss-RNA and hp-RNA were dissolved in TE buffer (10 mM Tris–HCl pH 7.5 and 1 mM EDTA) to achieve a sample concentration of 100 μM and used in the experiments at 4 μM final concentration. UPF2L was used at 10 μM concentration in a 200 μL volume of 20 mM Hepes pH 7.5, 150 mM NaF. The CD spectra were recorded over a 200-340 nm wavelength range at 20°C, with a 1 nm data interval, using a Jasco J-810 Spectropolarimeter, in 1 mm path-length cell. Scanning was performed at 50 nm/min and 8 accumulations were recorded for each spectrum. The buffer spectrum was subtracted from the sample spectra. The spectra of RNA only were further subtracted from the UPF2L / RNA spectra for analysis of RNA tertiary structure changes. Vice versa, the spectrum of UPF2L was subtracted from the UPF2L/ RNA spectra for analysis of protein structural changes. Analysis of spectra was performed using GraphPad Prism7 software.

### Fluorescence spectroscopy

Fluorescence spectroscopy was carried out using a CLARIOstar® Plus plate reader (BMG Labtech). The molecular beacon was dissolved in TE buffer (10 mM Tris–HCl pH 7.5 and 1 mM EDTA) to obtain a concentration of 100 μM, following the manufacturer recommendations. RNA samples at a concentration of 250 nM were incubated at 25 °C for 15min in a Corning® 384 well microplate (low volume black polystyrene, flat bottom). Fluorescence excitation and emission were recorded at 539 nm and 559 nm respectively. Subsequently, indicated concentrations of UPF2L were added in the wells, and the incubation was continued at 25°C for 1 h. Experiments were performed in duplicate in 20 μL final volume. The obtained data were plotted after subtracting the background fluorescence signals using GraphPad Prism7 software.

### Florescence anisotropy measurements

Fluorescence anisotropy analyses were performed using a Jobin Yvon Fluorology-3 (HORIBA Scientific) instrument. HEX-labelled oligonucleotides (Supplemental Table S1) were diluted to 10 nM in 150 µl assay buffer (25 mM HEPES pH 7.45, 150 mM NaCl, 5% glycerol) and placed in a 10 x 2 mm Quartz Suprasil cuvette (Hellma Analytics). Calibration and a zero-point reading were measured for excitation at 530 nm and emission at 550 nm. 1 µl UPF2L or MIF4G-D3 was added stepwise at different concentrations to achieve a serial dilution of protein within the cuvette. Measurements were performed using an integration time of 0.5 seconds over four accumulations and repeated in triplicate for each concentration. Assays were repeated in triplicate. Resulting signals were plotted using GraphPad Prism7 with the equation:

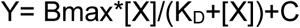

(referring to a one site specific binding (where Bmax is the total number of specific binding and X is the concentration of free protein) used to calculate the equilibrium dissociation constant (K_D_) and standard deviation).

### Sample preparation and negative-stain electron microscopy (EM)

Purified UPF2L in SEC buffer (25 mM HEPES pH 7.45, 150 nM NaCl, 1 mM TCEP, 2 mM MgCl2, 5% Glycerol) was used for negative-stain EM sample preparation. Complex formation was achieved by adding a 4-fold molar excess of RNA to UPF2L and incubating on ice for 1 hour. The sample was centrifuged at 16,200 x g for 10 mins and immediately used for grid preparation.

Carbon-coated copper 300 mesh grids (Electron Microscopy Sciences) were glow-discharged (15 sec at 15 mA) on a Leica EM ACE600 instrument (Leica Microsystems). 5 µl of UPF2L or UPF2L / RNA complex was applied to the grids at a concentration of 0.01 to 0.005 mg/ml and incubated for 1 min before blotting with Whatman filter paper (GE Healthcare). 5 µl of 3% (w/v) uranyl-acetate was applied and immediately blotted before applying an additional 5 µl of uranyl-acetate and incubating for 15-30 seconds followed by blotting. Data was collected using a FEI T20 200 kV Twin lens TEM microscope (within the Wolfson Bioimaging Suite, University of Bristol) and imaged at 68,000x magnification corresponding to a pixel size of 1.44 Å using an Ceta 4k x 4k charge-coupled device (CCD) camera (FEI).

### Negative-stain EM image processing

Images were processed using the RELION 3.1 software package (Scheres 2012). UPF2L micrographs were imported without contrast transfer function (CTF) correction. 2500 particles were manually picked and used for two rounds of 2D classifications using 25 classes. Eight classes were selected as templates for auto-picking, where 95,478 particles were picked. The auto-picked particles were subjected to multiple rounds of 2D classifications. A subset of class selections was used to generate final 2D class averages. The 2D classes of UPF2L were used as a template to auto pick 95,962 particles in micrographs with UP2L / hp-RNA sample. 224,197 particles were picked from UPF2L / ss-RNA micrographs. 2D class averages containing most particles are shown in Fig. 5 and Supplemental Fig. S8.

### RNA and protein structure predictions

The RNA 2D models were predicted using the Vienna RNA websuite (Lorenz et al. 2011) and MXfold2 (Sato et al. 2021). The RNA 3D models were predicted using 3dRNA/DNA (Zhang et al. 2022) and trRosettaRNA (Wang et al. 2023). To predict 2D and 3D dimer models of ss-RNA, a U_10_ or A_10_ linker sequence was inserted linking two ss-RNA molecules. This linker was then deleted from 2D and 3D models for visualization (Sato et al. 2021).

The UPF2L structure predictions were acquired from the ColabFold: AlphaFold2 Protein Structure Database (Mirdita et al. 2022). The five output predicted models were automatically ranked (1-best to 5-worst) based on highest average predicted local distance difference test (pLDDT) score for single structures and predicted template modelling (pTM) score for complexes. Subsequently, input UPF2L structures were relaxed to a low energy state according to backbone, relaxation, sidechain repacking and bond strength using the Relax application, as part of the Rosetta software suite (www.rosettacommons.org) (Tyka et al. 2011). Models were subjected to three cycles of relaxing to generate five models, ranked by total score, relating to the energy values from 19 tested terms including the attraction and repulsion energy between atoms, intermolecular bond forces (Van der Waals, hydrogen bond and disulphide bond) and side chain torsion angles (for full list of test terms see (Alford et al. 2017)).

## DATA DEPOSITION

All data for this manuscript are contained within the main article and Supplemental Information.

## SUPPLEMENTAL MATERIAL

Supplemental information comprises 7 figures and 1 table.

## ACKNOWLEDGEMENTS

We thank all past and present members of the Schaffitzel and Berger teams for their assistance and advice. We acknowledge support by the staff of the BrisSynBio BioSuite and the Wolfson Bioimaging Facility, in particular Judith Mantell. We thank Robert Barringer for help with the Relax Rosetta programme. C.S. acknowledges support by a Wellcome Trust Investigator award (210701/Z/18/Z). J.A.S. and C.S. acknowledge support by the Biotechnology and Biological Sciences Research Council-funded Southwest Biosciences Doctoral Training Partnership (DTP2: BB/M009122/1). M.V.V. and C.S. acknowledge funding from the European Union’s Horizon 2020 research and innovation programme under the Marie Sklodowska-Curie grant agreement (101024558). I.B. acknowledges support by the Wellcome Trust (106115/Z/14/Z). L.S. is funded by the Chinese Scholarship Council. For the purpose of Open Access, the authors have applied a CC BY public copyright license to any Author Accepted Manuscript version arising from this submission.

## REFERENCES

Alford RF, Leaver-Fay A, Jeliazkov JR, O’Meara MJ, DiMaio FP, Park H, Shapovalov MV, Renfrew PD, Mulligan VK, Kappel K et al. 2017. The Rosetta All-Atom Energy Function for Macromolecular Modeling and Design. J Chem Theory Comput 13: 3031–3048.

Buchwald G, Ebert J, Basquin C, Sauliere J, Jayachandran U, Bono F, Le Hir H, Conti E. 2010. Insights into the recruitment of the NMD machinery from the crystal structure of a core EJC-UPF3b complex. Proc Natl Acad Sci U S A 107: 10050–10055.

Bufton JC, Powers KT, Szeto JA, Toelzer C, Berger I, Schaffitzel C. 2022. Structures of nonsense-mediated mRNA decay factors UPF3B and UPF3A in complex with UPF2 reveal molecular basis for competitive binding and for neurodevelopmental disorder-causing mutation. Nucleic Acids Res 50: 5934–5947.

Bühler M, Steiner S, Mohn F, Paillusson A, Mühlemann O. 2006. EJC-independent degradation of nonsense immunoglobulin-mu mRNA depends on 3’ UTR length. Nat Struct Mol Biol 13: 462–464.

Chakrabarti S, Jayachandran U, Bonneau F, Fiorini F, Basquin C, Domcke S, Le Hir H, Conti E. 2011. Molecular mechanisms for the RNA-dependent ATPase activity of Upf1 and its regulation by Upf2. Mol Cell 41: 693–703.

Chamieh H, Ballut L, Bonneau F, Le Hir H. 2008. NMD factors UPF2 and UPF3 bridge UPF1 to the exon junction complex and stimulate its RNA helicase activity. Nat Struct Mol Biol 15: 85–93.

Chan WK, Bhalla AD, Le Hir H, Nguyen LS, Huang L, Gecz J, Wilkinson MF. 2009. A UPF3-mediated regulatory switch that maintains RNA surveillance. Nat Struct Mol Biol 16: 747–753.

Chapman JH, Craig JM, Wang CD, Gundlach JH, Neuman KC, Hogg JR. 2022. UPF1 mutants with intact ATPase but deficient helicase activities promote efficient nonsense-mediated mRNA decay. Nucleic Acids Res 50: 11876–11894.

Chapman JH, Youle AM, Grimme AL, Neuman KC, Hogg JR. 2024. UPF1 ATPase autoinhibition and activation modulate RNA binding kinetics and NMD efficiency. Nucleic Acids Res 52: 5376–5391.

Chauca-Diaz AM, Choi YJ, Resendiz MJ. 2015. Biophysical properties and thermal stability of oligonucleotides of RNA containing 7,8-dihydro-8-hydroxyadenosine. Biopolymers 103: 167–174.

Cheng Z, Muhlrad D, Lim MK, Parker R, Song H. 2007. Structural and functional insights into the human Upf1 helicase core. EMBO J 26: 253–264.

Chousal JN, Sohni A, Vitting-Seerup K, Cho K, Kim M, Tan K, Porse B, Wilkinson MF, Cook-Andersen H. 2022. Progression of the pluripotent epiblast depends upon the NMD factor UPF2. Development 149.

Clerici M, Deniaud A, Boehm V, Gehring NH, Schaffitzel C, Cusack S. 2014. Structural and functional analysis of the three MIF4G domains of nonsense-mediated decay factor UPF2. Nucleic Acids Res 42: 2673–2686.

Clerici M, Mourao A, Gutsche I, Gehring NH, Hentze MW, Kulozik A, Kadlec J, Sattler M, Cusack S. 2009. Unusual bipartite mode of interaction between the nonsense-mediated decay factors, UPF1 and UPF2. EMBO J 28: 2293–2306.

Deniaud A, Karuppasamy M, Bock T, Masiulis S, Huard K, Garzoni F, Kerschgens K, Hentze MW, Kulozik AE, Beck M et al. 2015. A network of SMG-8, SMG-9 and SMG-1 C-terminal insertion domain regulates UPF1 substrate recruitment and phosphorylation. Nucleic Acids Res 43: 7600–7611.

Fiorini F, Bagchi D, Le Hir H, Croquette V. 2015. Human Upf1 is a highly processive RNA helicase and translocase with RNP remodelling activities. Nat Commun 6: 7581.

Fritz SE, Ranganathan S, Wang CD, Hogg JR. 2022. An alternative UPF1 isoform drives conditional remodeling of nonsense-mediated mRNA decay. EMBO J 41: e108898.

Gehring NH, Hentze MW, Kulozik AE. 2008. Tethering assays to investigate nonsense-mediated mRNA decay activating proteins. Methods Enzymol 448: 467–482.

Gehring NH, Kunz JB, Neu-Yilik G, Breit S, Viegas MH, Hentze MW, Kulozik AE. 2005. Exon-junction complex components specify distinct routes of nonsense-mediated mRNA decay with differential cofactor requirements. Mol Cell 20: 65–75.

Gehring NH, Lamprinaki S, Hentze MW, Kulozik AE. 2009. The hierarchy of exon-junction complex assembly by the spliceosome explains key features of mammalian nonsense-mediated mRNA decay. PLoS Biol 7: e1000120.

Gupta K, Tolzer C, Sari-Ak D, Fitzgerald DJ, Schaffitzel C, Berger I. 2019. MultiBac: Baculovirus-Mediated Multigene DNA Cargo Delivery in Insect and Mammalian Cells. Viruses 11.

Hauer C, Sieber J, Schwarzl T, Hollerer I, Curk T, Alleaume AM, Hentze MW, Kulozik AE. 2016. Exon Junction Complexes Show a Distributional Bias toward Alternatively Spliced mRNAs and against mRNAs Coding for Ribosomal Proteins. Cell Rep 16: 1588–1603.

Hogg JR, Goff SP. 2010. Upf1 senses 3’UTR length to potentiate mRNA decay. Cell 143: 379–389.

Hwang H, Myong S. 2014. Protein induced fluorescence enhancement (PIFE) for probing protein-nucleic acid interactions. Chem Soc Rev 43: 1221–1229.

Jaffrey SR, Wilkinson MF. 2018. Nonsense-mediated RNA decay in the brain: emerging modulator of neural development and disease. Nat Rev Neurosci 19: 715–728.

Kadlec J, Izaurralde E, Cusack S. 2004. The structural basis for the interaction between nonsense-mediated mRNA decay factors UPF2 and UPF3. Nat Struct Mol Biol 11: 330–337.

Karousis ED, Mühlemann O. 2019. Nonsense-Mediated mRNA Decay Begins Where Translation Ends. Cold Spring Harb Perspect Biol 11.

Kashima I, Yamashita A, Izumi N, Kataoka N, Morishita R, Hoshino S, Ohno M, Dreyfuss G, Ohno S. 2006. Binding of a novel SMG-1-Upf1-eRF1-eRF3 complex (SURF) to the exon junction complex triggers Upf1 phosphorylation and nonsense-mediated mRNA decay. Genes Dev 20: 355–367.

Kurosaki T, Popp MW, Maquat LE. 2019. Quality and quantity control of gene expression by nonsense-mediated mRNA decay. Nat Rev Mol Cell Biol 20: 406–420.

Lavysh D, Neu-Yilik G. 2020. UPF1-Mediated RNA Decay-Danse Macabre in a Cloud. Biomolecules 10.

Le Hir H, Sauliere J, Wang Z. 2016. The exon junction complex as a node of post-transcriptional networks. Nat Rev Mol Cell Biol 17: 41–54.

Lee SR, Pratt GA, Martinez FJ, Yeo GW, Lykke-Andersen J. 2015. Target Discrimination in Nonsense-Mediated mRNA Decay Requires Upf1 ATPase Activity. Mol Cell 59: 413–425.

Lejeune F. 2022. Nonsense-Mediated mRNA Decay, a Finely Regulated Mechanism. Biomedicines 10.

Lindeboom RG, Supek F, Lehner B. 2016. The rules and impact of nonsense-mediated mRNA decay in human cancers. Nat Genet 48: 1112–1118.

Lopez-Perrote A, Castano R, Melero R, Zamarro T, Kurosawa H, Ohnishi T, Uchiyama A, Aoyagi K, Buchwald G, Kataoka N et al. 2016. Human nonsense-mediated mRNA decay factor UPF2 interacts directly with eRF3 and the SURF complex. Nucleic Acids Res 44: 1909–1923.

Lorenz R, Bernhart SH, Honer Zu Siederdissen C, Tafer H, Flamm C, Stadler PF, Hofacker IL. 2011. ViennaRNA Package 2.0. Algorithms Mol Biol 6: 26.

Lykke-Andersen J, Shu MD, Steitz JA. 2000. Human Upf proteins target an mRNA for nonsense-mediated decay when bound downstream of a termination codon. Cell 103: 1121–1131.

Mabin JW, Woodward LA, Patton RD, Yi Z, Jia M, Wysocki VH, Bundschuh R, Singh G. 2018. The Exon Junction Complex Undergoes a Compositional Switch that Alters mRNP Structure and Nonsense-Mediated mRNA Decay Activity. Cell Rep 25: 2431–2446 e2437.

Melero R, Buchwald G, Castano R, Raabe M, Gil D, Lazaro M, Urlaub H, Conti E, Llorca O. 2012. The cryo-EM structure of the UPF-EJC complex shows UPF1 poised toward the RNA 3’ end. Nat Struct Mol Biol 19: 498–505, S491-492.

Metze S, Herzog VA, Ruepp MD, Mühlemann O. 2013. Comparison of EJC-enhanced and EJC-independent NMD in human cells reveals two partially redundant degradation pathways. RNA 19: 1432–1448.

Mirdita M, Schutze K, Moriwaki Y, Heo L, Ovchinnikov S, Steinegger M. 2022. ColabFold: making protein folding accessible to all. Nat Methods 19: 679–682.

Mort M, Ivanov D, Cooper DN, Chuzhanova NA. 2008. A meta-analysis of nonsense mutations causing human genetic disease. Hum Mutat 29: 1037–1047.

Nasif S, Contu L, Mühlemann O. 2018. Beyond quality control: The role of nonsense-mediated mRNA decay (NMD) in regulating gene expression. Semin Cell Dev Biol 75: 78–87.

Neu-Yilik G, Raimondeau E, Eliseev B, Yeramala L, Amthor B, Deniaud A, Huard K, Kerschgens K, Hentze MW, Schaffitzel C et al. 2017. Dual function of UPF3B in early and late translation termination. EMBO J 36: 2968–2986.

Rajkowitsch L, Chen D, Stampfl S, Semrad K, Waldsich C, Mayer O, Jantsch MF, Konrat R, Blasi U, Schroeder R. 2007. RNA chaperones, RNA annealers and RNA helicases. RNA Biol 4: 118–130.

Sato K, Akiyama M, Sakakibara Y. 2021. RNA secondary structure prediction using deep learning with thermodynamic integration. Nat Commun 12: 941.

Scheres SH. 2012. RELION: implementation of a Bayesian approach to cryo-EM structure determination. J Struct Biol 180: 519–530.

Schlautmann LP, Gehring NH. 2020. A Day in the Life of the Exon Junction Complex. Biomolecules 10.

Serdar LD, Whiteside DL, Baker KE. 2016. ATP hydrolysis by UPF1 is required for efficient translation termination at premature stop codons. Nat Commun 7: 14021.

Serdar LD, Whiteside DL, Nock SL, McGrath D, Baker KE. 2020. Inhibition of post-termination ribosome recycling at premature termination codons in UPF1 ATPase mutants. Elife 9.

Szpotkowski K, Wojcik K, Kurzynska-Kokorniak A. 2023. Structural studies of protein-nucleic acid complexes: A brief overview of the selected techniques. Comput Struct Biotechnol J 21: 2858–2872.

Tan K, Stupack DG, Wilkinson MF. 2022. Nonsense-mediated RNA decay: an emerging modulator of malignancy. Nat Rev Cancer 22: 437–451.

Tyagi S, Kramer FR. 1996. Molecular beacons: probes that fluoresce upon hybridization. Nat Biotechnol 14: 303–308.

Tyka MD, Keedy DA, Andre I, Dimaio F, Song Y, Richardson DC, Richardson JS, Baker D. 2011. Alternate states of proteins revealed by detailed energy landscape mapping. J Mol Biol 405: 607–618.

Wang W, Feng C, Han R, Wang Z, Ye L, Du Z, Wei H, Zhang F, Peng Z, Yang J. 2023. trRosettaRNA: automated prediction of RNA 3D structure with transformer network. Nat Commun 14: 7266.

Xue G, Maciej VD, Machado de Amorim A, Pak M, Jayachandran U, Chakrabarti S. 2023. Modulation of RNA-binding properties of the RNA helicase UPF1 by its activator UPF2. RNA 29: 178–187.

Zhang Y, Wang J, Xiao Y. 2022. 3dRNA: 3D Structure Prediction from Linear to Circular RNAs. J Mol Biol 434: 167452.

